# Massively parallel profiling of HIV-1 resistance to the fusion inhibitor enfuvirtide

**DOI:** 10.1101/472746

**Authors:** Adam S. Dingens, Dana Arenz, Julie Overbaugh, Jesse D. Bloom

## Abstract

Resistance to enfuvirtide, the only clinically approved HIV-1 entry inhibitor, has primarily been mapped to the binding site in the N-terminal heptad repeat (NHR) of the Env transmembrane domain and a limited number of allosteric sites. To better delineate the genotypic determinants of resistance, we used deep mutational scanning to quantify how all mutations to HIV-1 Env affect enfuvirtide sensitivity. We identified numerous additional resistance mutations in the NHR and other regions of Env, including the co-receptor binding site. This complete map of resistance sheds light on the diverse mechanisms of enfuvirtide resistance and can inform clinical monitoring of patients.

## Introduction

Antiretroviral drug therapy has reduced the global burden of HIV/AIDS. However, HIV-1’s exceptional evolutionary capacity enables the virus to evolve resistance, eroding the therapeutic efficacy of drug regimens. Identifying drug resistance mutations is therefore imperative to the clinical management of HIV-1/AIDS, particularly as uptake of antiviral treatment is increasing globally.

Enfuvirtide (also known as T-20 and Fuzeon), the only clinically approved fusion inhibitor, is a 36 amino-acid peptide derived from the C-terminal Heptad repeat (CHR) domain of gp41 that inhibits viral entry into cells. After Env binds CD4, enfuvirtide binds to the coiled-coil NHR structure of the pre-hairpin intermediate and inhibits fusion by preventing the CHR from binding to the NHR to form the 6-helical bundle [1]. Due to enfuvirtide’s high cost and need for twice-daily subcutaneous injections, it is used as part of a “salvage therapy” for highly treatment-experienced patients with multidrug resistance. Such patients need to be carefully monitored for resistance before and during therapy [2], which requires knowing the viral genotypic determinants of enfuvirtide resistance.

Prior studies that analyzed enfuvirtide resistance in cell culture [3] and *in vivo* during therapy [4, 5] identified resistance mutations at sites 547-556, implicating this region of the NHR as enfuvirtide’s binding site. However, mutations in gp120 outside of this region have also been shown to affect enfuvirtide sensitivity, possibly by altering co-receptor tropism, co-receptor affinity, or fusion kinetics [6, 7].

Here we more completely map resistance mutations by using deep mutational scanning to quantify how enfuvirtide resistance is affected by all mutations to HIV-1 Env compatible with viral replication.

## Methods

### Generation of Env mutant virus libraries

We used Env mutant virus libraries from a subtype A transmitted variant BG505 to map enfuvirtide resistance. The generation of these libraries has been previously described [8, 9]. Briefly, we independently generated triplicate proviral DNA libraries encoding codon-level mutations to sites 31-702 (HXB2 numbering is used throughout the manuscript) of BG505.W6M.C2.T332N *env.* These libraries contain (670 mutagenized sites) × (19 amino acid mutations) = 12,730 possible amino acid mutations. We then produced full-length viruses from these mutant DNA libraries and passaged them in SupT1.CCR5 cells to select for functional viruses, resulting in mutant virus libraries that encode all functionally tolerated mutations to Env [8].

### Resistance profiling

To identify resistance mutations, we incubated the mutant virus libraries with or without enfuvirtide, infected cells, and then identified the mutant viruses that were enriched upon drug selection using deep sequencing. This approach is similar to the mutational antigenic profiling process we have previously used to map antibody escape [10, 11]. Briefly, 5×10^5^ to 1×10^6^ infectious units of three independent mutant virus libraries were incubated in the presence of 8ug/mL of enfuvirtide, then infected into 1×10^6^ SupT1.CCR5 cells in R10 (RPMI with 10% FBS, 1% 200 mM L-glutamine, and 100 units/mL of penicillin and streptomycin), containing 100ug/mL DEAE-dextran. Three hours post infection, cells were pelleted and resuspended in 1 mL R10 without DEAE-dextran, and at 12 hours post infection, non-integrated viral cDNA was isolated using a miniprep. As mock-selected controls, each mutant virus library was infected into cells without enfuvirtide selection, as were four 10-fold serial dilutions of each mutant virus library. Selected and mock-selected viral cDNA was then sequenced with a barcoded subamplicon sequencing approach as previously described [8], which introduces unique molecular identifiers used to correct sequencing errors. The amount of virus library that entered cells was quantified via qPCR [11].

### Analysis of deep sequencing data

We used dms_tools2 version 2.3.0 (https://jbloomlab.github.io/dmstools2/) to analyze the deep sequencing data [12]. The differential selection statistic has been previously described [13] and is documented at https://jbloomlab.github.io/dmstools2/diffsel.html. Sequencing of wildtype proviral DNA plasmid was used as the error control during the calculation of the differential selection.

### Data availability and source code

The entire computational analysis is provided as a Jupyter Notebook (File S1) and at https://github.com/jbloomlab/EnfuvirtideResistance. Differential selection measurements are provided as File S2. Illumina deep sequencing reads were deposited into the NCBI SRA as SRR8097918-SRR8097920.

### TZM-bl inhibition assays

Individual mutations were introduced into BG505.T332N Env, and pseuodviruses were generated with these Envs and tested in TZM-bl neutralization assays using previously described methods.

## Results

To quantify the effect of all mutations to Env on enfuvirtide sensitivity, we selected triplicate BG505 mutant virus libraries with 8 ug/mL enfuvirtide. This highly selective drug concentration, which resulted in just 0.15% to 0.78% of the mutant virus libraries surviving selection, was used to mimic the environment in which resistance could arise during therapy. We deep sequenced the *env* genes of viruses that entered cells and calculated the enrichment of each mutation in the drug-selected condition relative to a non-selected mutant virus library, to calculate a quantity that we term the *differential selection* [13].

Selection of resistance mutations was highly reproducible across biological triplicates (Supplementary Figure 1). Most resistance mutations were found in the NHR domain (Figure 1, Supplementary Figure 2) and included many well-characterized mutations on the IAS–USA drug resistance mutations list [14].

**Figure 1.**
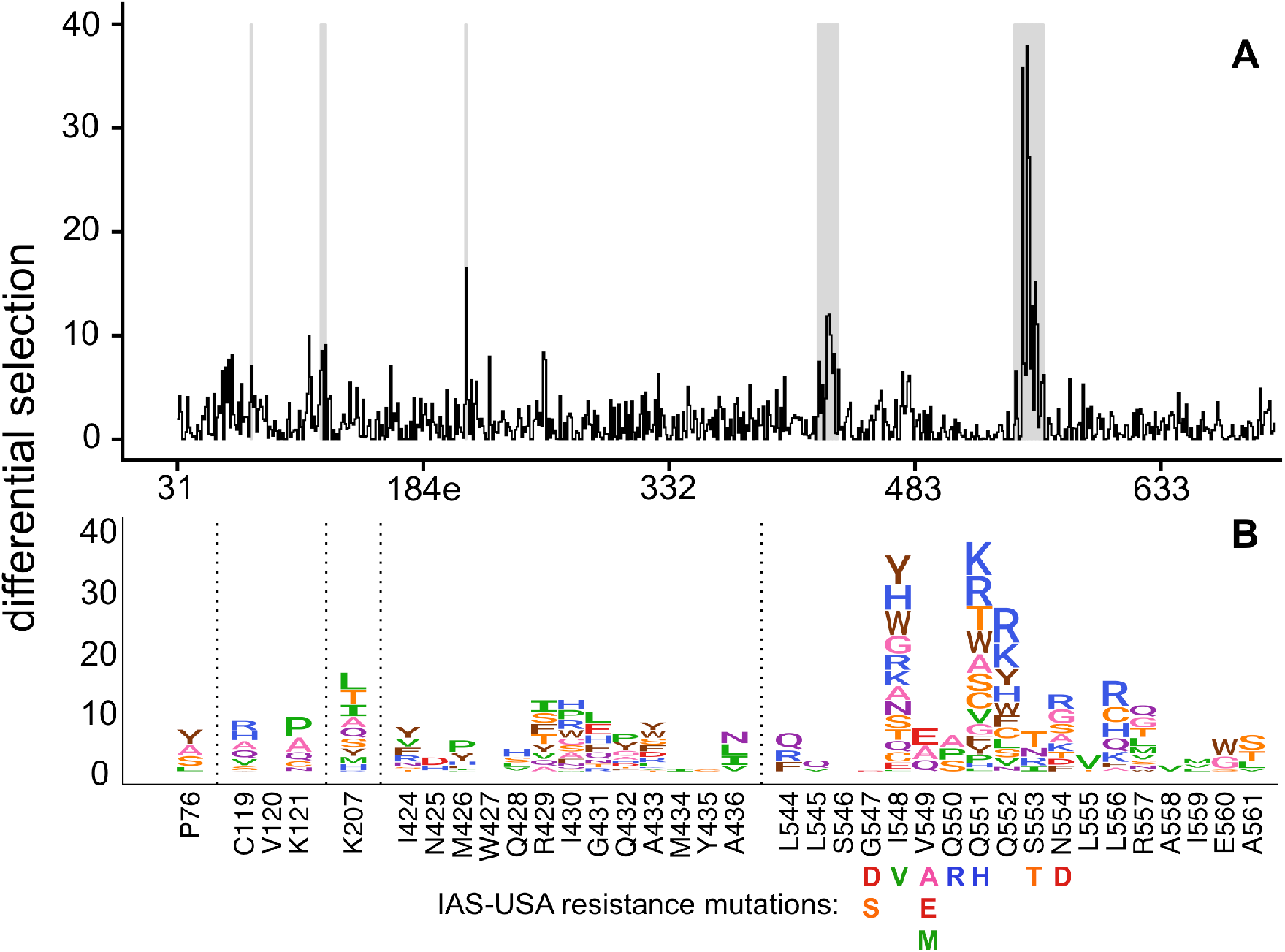
Complete map of enfuvirtide resistance. **A**. The positive site differential selection is plotted across the mutagenized portion of Env. **B**. The mutation-level resistance profile for regions of interest (highlighted in grey in A). The height of each amino acid is proportional to its differential selection. All mutations from the 2017 IAS-USA enfuvirtide resistance mutations list [14] are labeled below their site.

For example, S553T, a known resistance mutation, was the largest effect size mutation at site 553 in our experiments. Similarly, V549E and V549A were the largest effect size mutations at site 549 in our experiments, and both have been previously shown to cause resistance. However, V549M was not enriched in our experiments despite being associated with resistance in other strains. Similarly, the known I548V resistance mutation was not enriched in our experiments, but numerous other mutations at site 548 were strongly enriched. These discrepancies may be due to strain specific differences in enfuvirtide sensitivity, discussed in more detail in the Discussion.

We also identified resistance mutations in the NHR region that had not been previously characterized. For example, while Q551H is a known resistance mutation, our data revealed that many additional amino acids at this site also increased enfuvirtide resistance, with 12 other mutations at this site having a larger effect than Q551H (Figure 1). Further, mutations to additional sites in NHR not included in the IAS-USA drug resistance mutations list, such as sites 552, 556, 557, and 560, also increased resistance to enfuvirtide (Figure 1).

Many of these resistance mutations cluster in the binding site of enfuvirtide in the NHR (Figure 2A). Site 551, where many different mutations confer resistance, directly interacts with enfuvirtide. Surprisingly, the side chains at the sites of the largest-effect mutations (548 and 552) face other NHR α-helices or the NHR trimer center (Figure 2A), and resistance at these sites occurred primarily due to mutations to positively charged or bulky amino acids (Figure 1B). The next two largest effect sites in this region, 554 and 556, appear to interact with enfuvirtide (Figure 2A).

**Figure 2.**
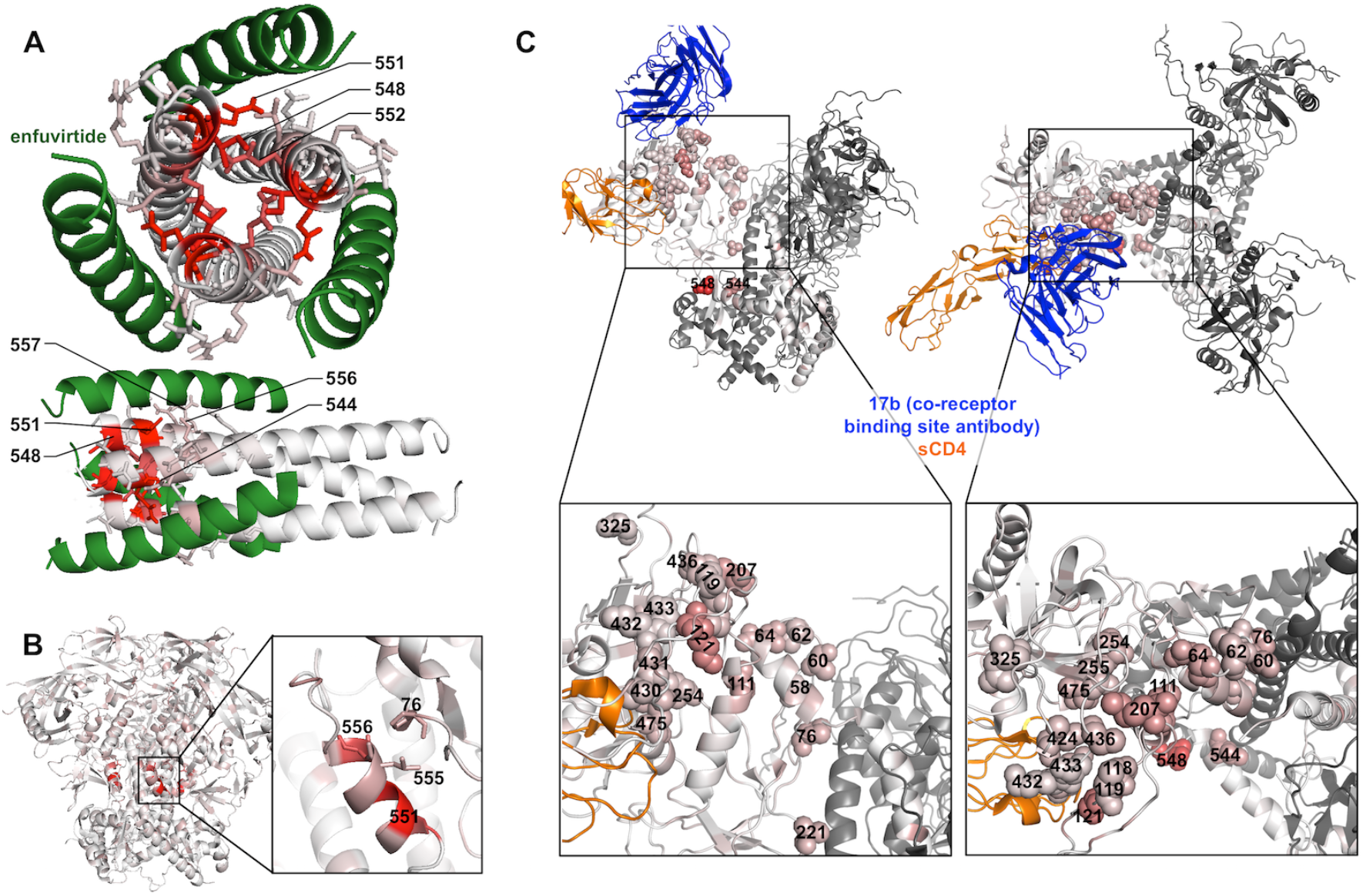
Locations of resistance mutations in different Env conformations. **A**. A model of enfuvirtide (green) binding to the NHR coiled-coil trimer (colored according to the positive site differential selection at each site). The top 5% of all resistance sites are shown with sticks. While there is not a structure of enfuvirtide bound to Env, we used the post fusion, 6-helical bundle structure, composed of NHR and CHR peptides, to model enfuvirtide binding. This model assumes that enfuvirtide binding to NHR is similar to the CHR peptide binding to NHR in this structure. Portions of the CHR present in the structure that do not overlap with enfuvirtide are omitted, and the entirety of enfuvirtide is not present in the structure. PDB: 1AIK **B**. The closed, pre-fusion conformation of BG505 Env colored according to the maximum mutation differential at each site. PDB: 5YFK **C**. The side-and top-view of post-CD4-bound, open structure of BG505 Env, also bound by the co-receptor binding site antibody 17b, colored according to the maximum mutation differential at each site. The top 5% of resistance sites are shown with spheres. PDB: 5VN3.

There was also modest but reproducible enrichment of mutations at other Env sites outside of the NHR domain. One such mutation was P76Y, which interacts with NHR sites L555 and L556 near the enfuvirtide binding site in the prefusion conformation (Figure 2B). Other resistance mutations occurred at sites 424-436 in the β20/β21 strand of C4, as well as sites 119, 121, and 207 in the V1/V2 stem. While the V1/V2 stem is distant from β20/β21 in the prefusion Env conformation, it shifts upon CD4 binding to form the 4-stranded bridging sheet along with the β20/β21 strand to create the co-receptor binding site (Figure 2C). Similarly, additional sites of modest resistance mutations, including sites 60, 62, 64, 111, 254, 255, and 475 all cluster near the co-receptor binding site in the open, CD4-bound state (Figure 2C).

To validate our high-throughput mapping of enfuvirtide resistance mutations, we generated and tested individual BG505 Env pseudoviruses bearing single mutations for enfuvirtide sensitivity. In the NHR domain, both the previously characterized V549E mutation and the Q552R mutation we identified increased resistance, shifting the IC_50_ by >150-fold (Supplementary Figure 3). Other mutations that were modestly enriched (P76Y, C119R, K121P, and K207L) had little effect on IC_50_ but instead decreased the maximal inhibition plateau at high concentrations of drug, including the 8 ug/mL concentration used in our experiments (Supplementary Figure 3). Notably, both these validation experiments and the high-throughput mapping were performed with 100ug/mL DEAE-dextran. When the TZM-bl assays were repeated with just 10ug/mL DEAE-dextran, some of the resistance phenotypes were not as prominent (Supplementary Figure 4).

## Discussion

We have quantified the effect of all single amino acid mutations to the extracellular and transmembrane ectodomain of BG505 Env on resistance to the fusion inhibitor enfuvirtide. This map of resistance mutations includes both previously characterized and numerous novel resistance mutations. This dataset may be of use in the clinical monitoring of resistance during therapy and the genotypic prediction of enfuvirtide sensitivity prior to treatment.

The selected mutations also help elucidate Env’s diverse mechanisms of enfuvirtide resistance, which likely include both disrupting enfuvirtide binding and altering fusion kinetics/conformations in multiple ways. While some mutations to NHR may directly disrupt interactions with enfuvirtide (e.g. site 551), other resistance mutations in this region appear to introduce positive charges or bulky amino acids at the center of the NHR coiled-coil. These mutations may slightly alter the coiled-coil structure to favor the intramolecular binding of the CHR domain over binding to enfuvirtide. It is possible that this dynamic is influenced by the subtype of the virus, as enfuvirtide is a mimetic of a subtype B CHR domain and differs from BG505 at 7 of 36 amino acids.

Other resistance mutations likely alter fusion kinetics or accessibility of the enfuvirtide binding site. P76Y may result in additional hydrophobic interactions between site 76 and L555 or L556 in the NHR in the prefusion conformation (Figure 2B). This interaction could directly limit access of enfuvirtide to the NHR binding site or delay the formation of the enfuvirtide-sensitive, CD4-induced conformational intermediate present post CD4-binding.

We also uncovered many small-effect size resistance mutations throughout the co-receptor binding site. It has been previously shown that co-receptor tropism, CCR5 affinity, and variability in V3 can affect enfuvirtide sensitivity [6, 7], but we characterized resistance mutations at numerous additional sites in or near the co-receptor binding site, including sites in the V1/V2 loop stem and C4. Such mutations may increase CCR5 receptor affinity and/or fusion kinetics, limiting the time in which a CD4-induced, enfuvirtide sensitive conformational intermediate is exposed. However, there is also evidence of weak interactions between enfuvirtide and the co-receptor binding site on gp120 [15]; resistance mutations in this region may disrupt this potential secondary enfuvirtide-Env interaction.

The complex mechanisms of enfuvirtide resistance were highlighted further when validating these resistance mutations with traditional inhibition curves. While mutations in the NHR domain resulted in a shift in the inhibition curve, the remaining mutants we tested resulted in a decrease in the maximum percent inhibition rather than a shift in the inhibition curve. It remains to be determined if such phenotypes manifest *in vivo.* However, if a small subpopulation of viruses were resistant to enfuvirtide at *in vivo* drug concentrations, this could present an opportunity for additional resistance to enfuvirtide or other drugs in combination therapies to arise.

While the resistance mutations uncovered here should be considered when evaluating clinical resistance, they also come with some caveats. We examined the effect of single amino acids in a single subtype A Env. Many patients receiving enfuvirtide have subtype B infections, and there could be strain-specific differences in the effects of mutations on enfuvirtide resistance. For example, we do not identify the G547D and G547S resistance mutations, which were identified by enfuvirtide selection of lab-adapted, subtype B viruses in cell culture [3]. Our previous studies show that these mutations are tolerated in a lab-adapted subtype B virus, but they are not well-tolerated by the transmitted subtype A variant BG505 during viral replication in cell culture [8, 9]. These differences highlight how the balance between enfuvirtide resistance and replicative fitness may depend on the virus strain.

In summary, this map of enfuvirtide resistance identifies numerous previously uncharacterized and potentially clinically relevant resistance mutations, highlighting the utility of using viral deep mutational scanning approaches to evaluate drug resistance. This data also sheds light on the mechanisms of enfuvirtide resistance and Env’s fusion process.

## Supplementary Figures

**Supplementary Figure 1.**
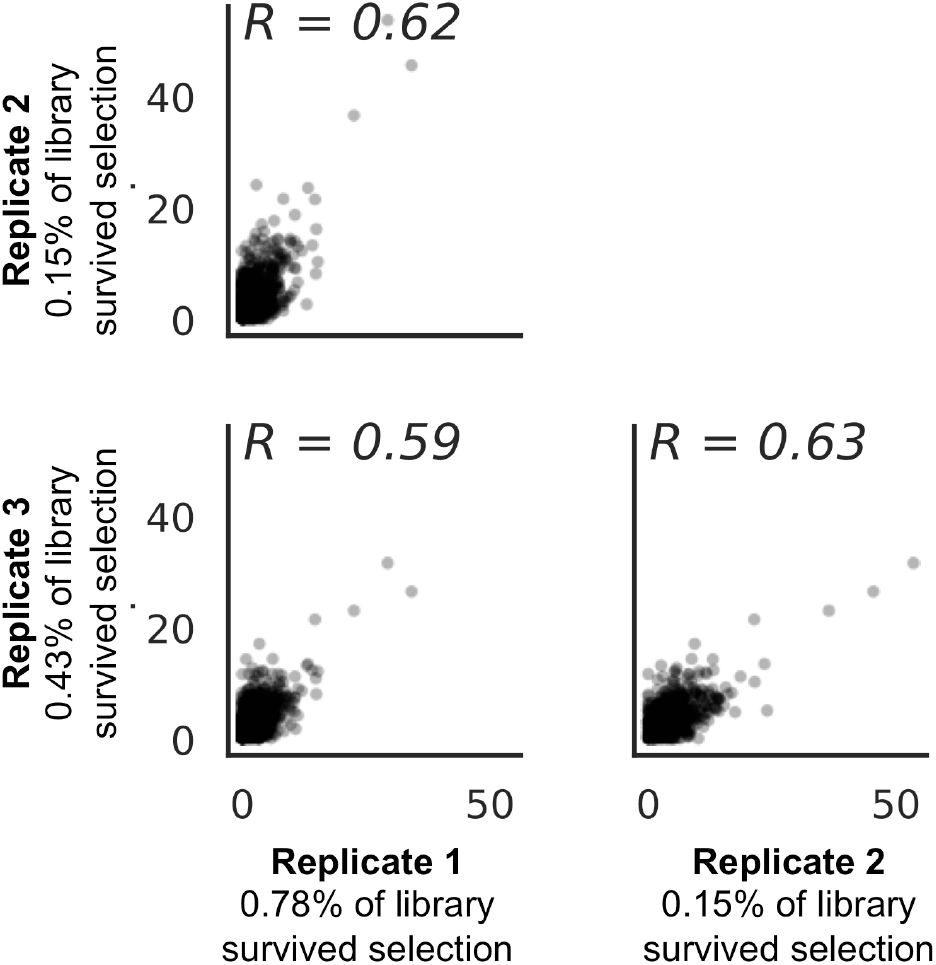
The correlations between the positive site differential selection for biological triplicates. The fraction of the virus that survived enfuvirtide selection, as calculated from a qPCR standard curve, is labeled for each replicate.

**Supplementary Figure 2.**
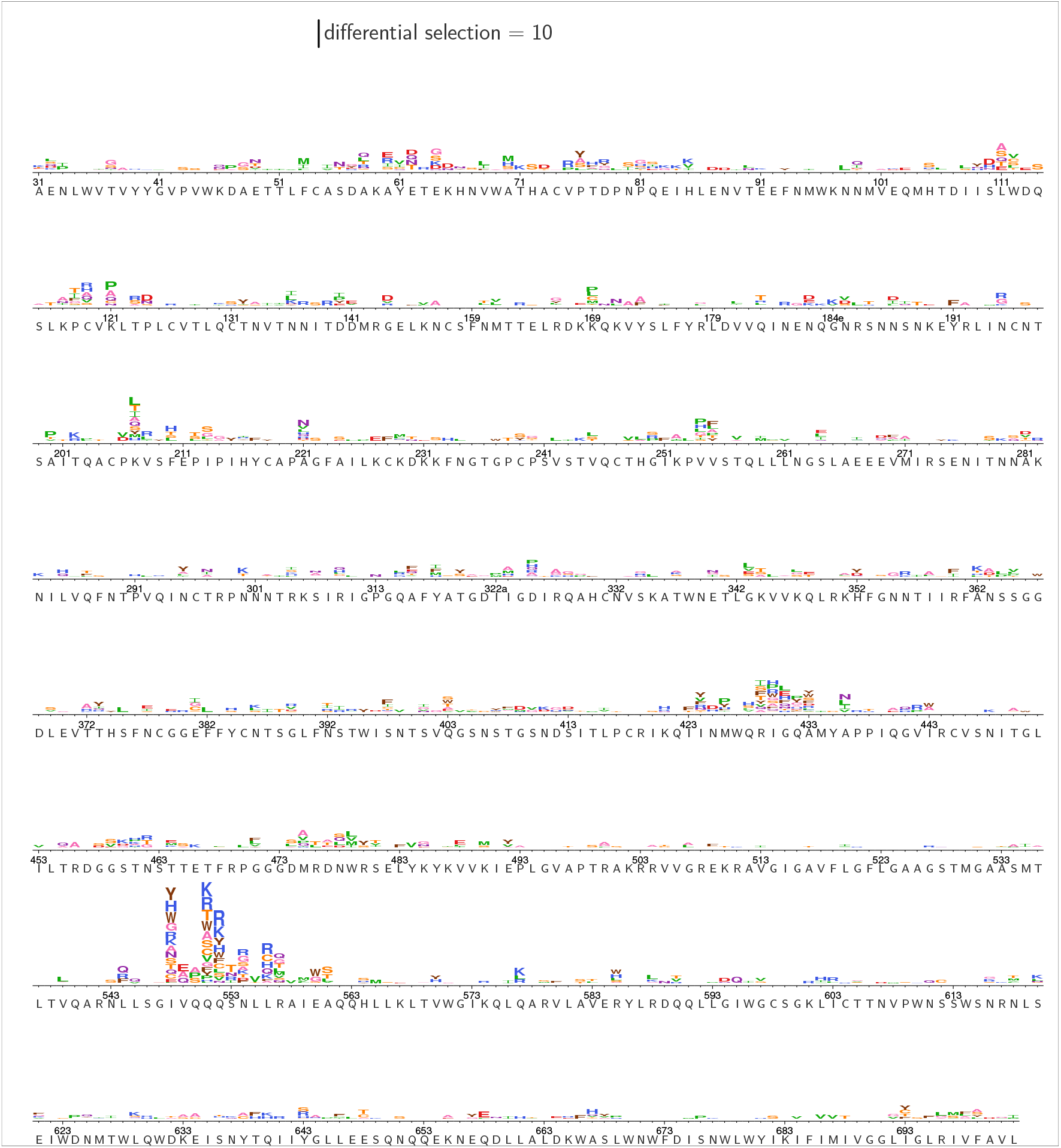
The complete, mutation-level resistance profile across Env. The height of each amino acid is proportional to its differential selection.

**Supplementary Figure 3.**
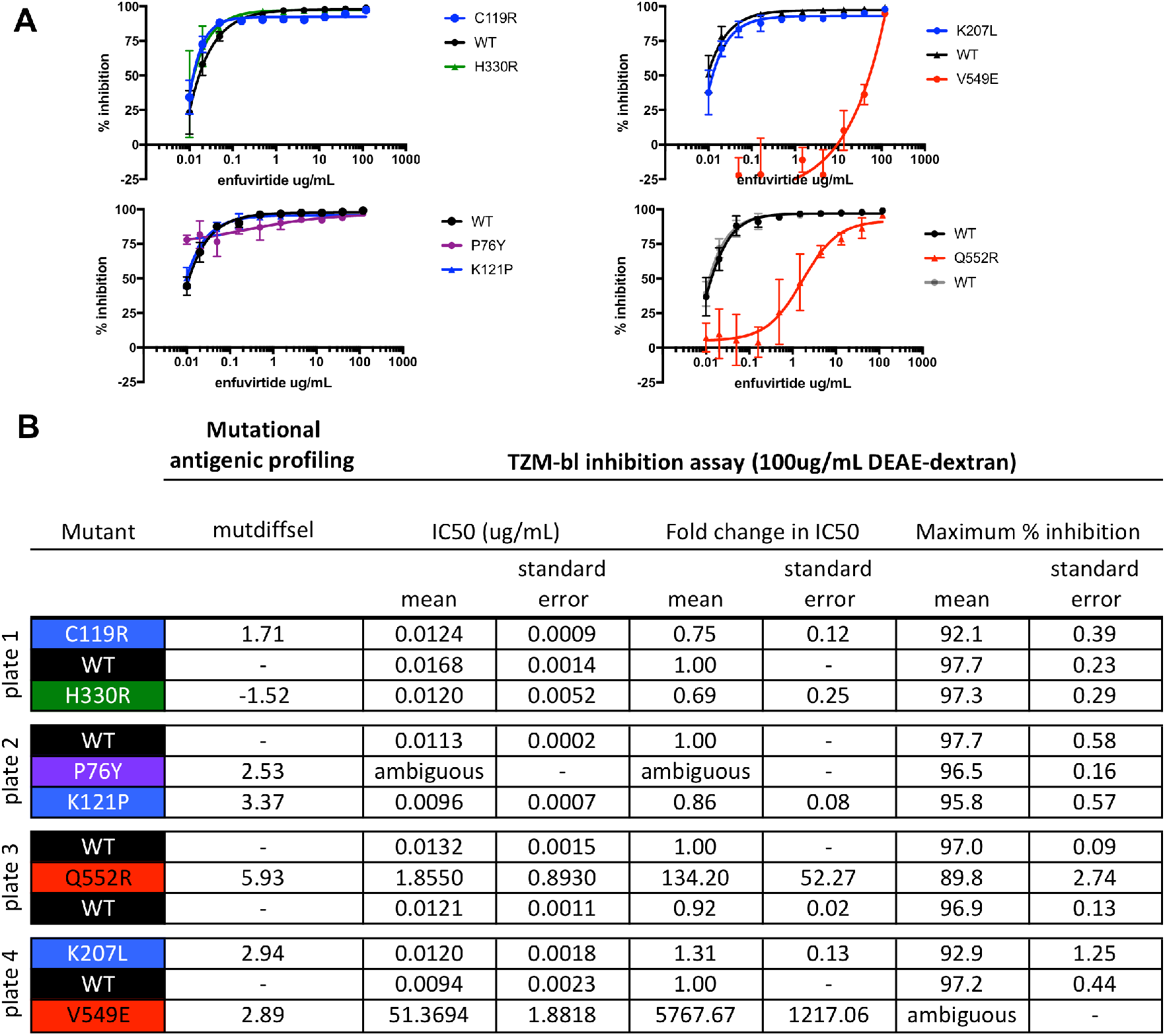
Validation of enfuvirtide resistance mutants using a TZM-bl inhibition assay. TZM-bl inhibition assays were performed in the presence of 100ug/mL DEAE-dextran, similar to the resistance profiling. **A**. Inhibition curves are the average of two biological replicates, each performed in duplicate. **B**. The IC_50_, the fold change in IC_50_ relative to wildtype (WT), and the maximum percent inhibition for each mutant, determined from the fit four-parameter logistic curves. At least one WT virus was run on each plate and each mutant virus curve was compared to the plate internal WT control. Values were computed independently for each experimental replicate, then averaged across replicates. The standard error of the mean is also shown. H330R, which was not enriched in the resistance profiling, was included as a control. In A and B, mutant psuedoviruses are colored according to groups (black: WT; green: control mutant not expected to affect enfuvirtide sensitivity; blue: mutants in the V1/V2 Stem/co-receptor binding site; red: mutants in/near NHR binding site).

**Supplementary Figure 4.**
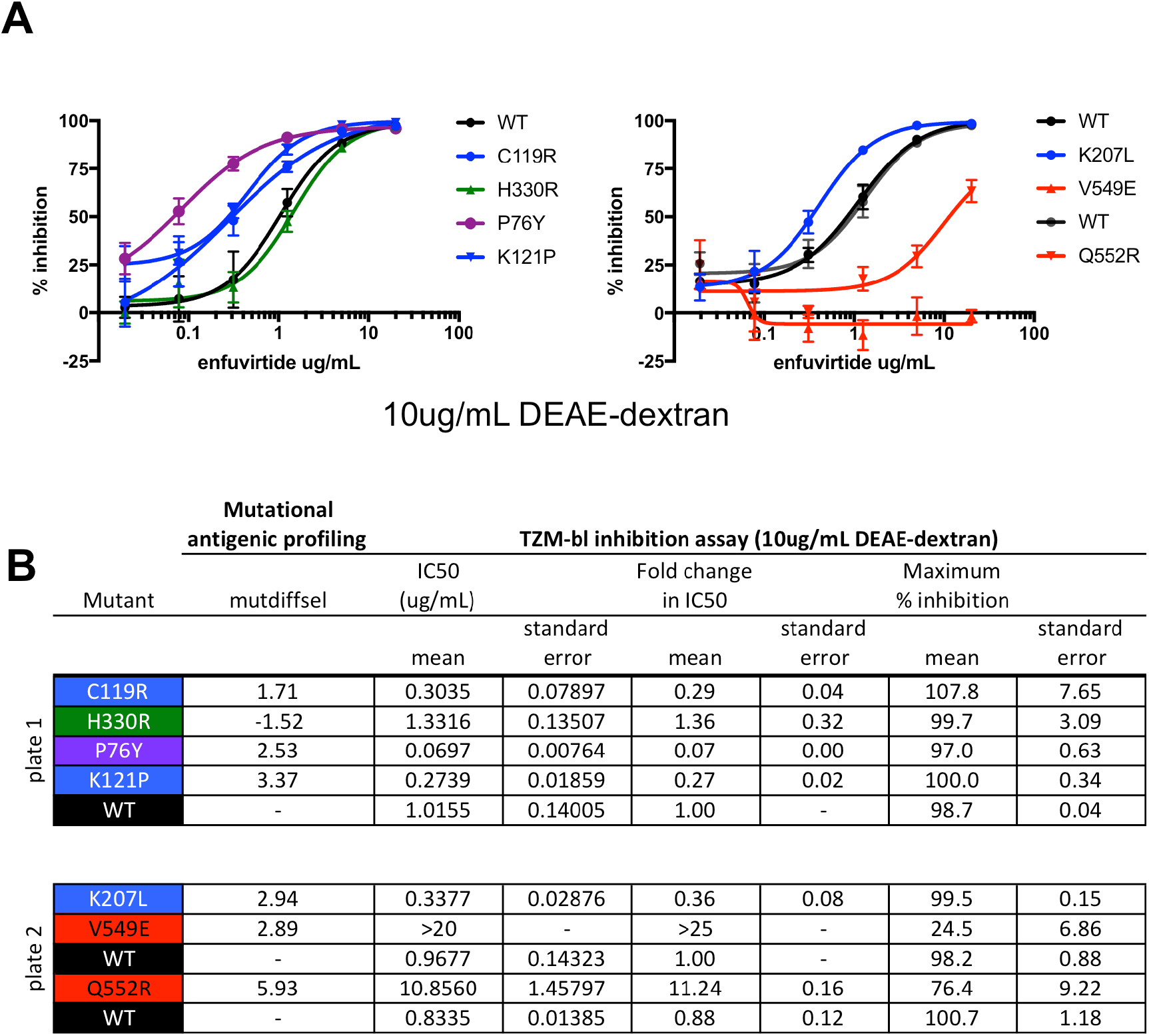
TZM-bl inhibition assay performed with 10ug/ml_ DEAE-dextran instead of the 100ug/mL concentration used in Supplementary Figure 3 and resistance profiling experiments. **A**. and **B**. As in Supplementary Figure 3.

## Supplementary File Legends

**Supplementary File 1**. An iPython notebook containing the entire computational analysis.

**Supplementary File 2**. A csv file containing the median differential selection estimates for enfuvirtide.

## Funding

ASD was supported by an NSF Graduate Research Fellowship [DGE-1256082]. This work was supported by the National Institutes of Health [R01AI127893 to JDB and DA039543 to JO]. JDB is an Investigator of the Howard Hughes Medical Institute. The following reagent was obtained through the NIH AIDS Reagent Program, Division of AIDS, NIAID, NIH: T-20, (Enfuvirtide).

## Acknowledgements

We thank Caelan Radford and Haidyn Weight for assistance with generating pseudoviruses and performing TZM-bl inhibition assays.

## Conflicts of interest

The authors have no conflicts of interest

## Author Contributions

Conceptualization, A.S.D.; Methodology, A.S.D., and J.D.B.; Validation, A.S.D. D. A.; Investigation, A.S.D.; Software, A.S.D.; Supervision, J.O. and J.D.B.; Writing – Original Draft, A.S.D.; Writing – Review & Editing, A.S.D., J.O., and J.D.B.; Funding Acquisition, J.O. and J.D.B.

